# Physiological Levels of Estrogen and Progesterone Modulate CR3 expression on, and *Neisseria gonorrhoeae* Infection of, Primary, Human, Cervical Epithelial Cells

**DOI:** 10.1101/2022.06.03.494700

**Authors:** Trevor J. Edwards, Jennifer L. Edwards

## Abstract

Despite advances made in our understanding of *Neisseria gonorrhoeae* pathogenesis, factors dictating the divergent presentation of gonococcal disease observed between men and women, *in vivo*, remain unclear. Clinical data indicate that gonococcal pathogenesis of the female genital tract is influenced by steroid hormones. Notwithstanding, there are limited data addressing how steroid hormones modulate gonococcal pathogenesis. Hence, we investigated the effect(s) of physiological concentrations of estrogens and progestogens on *N. gonorrhoeae* viability and on complement-mediated infection of primary cervical cells. In contrast to previous studies that showed a bacteriostatic effect of non-physiological concentrations of steroid hormones on gonococci, our data indicate that physiological concentrations of estrogens and progestogens do not inhibit gonococcal growth *in vitro* or during infection of primary cervical cells. Estradiol promoted complement receptor 3 recruitment to the cervical cell surface and, thus, the ability of gonococci to associate with these cells. Progesterone-predominant assay conditions resulted in decreased expression of Opa proteins by gonococci, increased complement production by cervical cells, and increased iC3b opsonization of gonococci during cervical cell challenge. Collectively, our data support clinical observations and demonstrate that estrogens and progestogens distinctly modulate gonococcal cervical infection.

## Introduction

*Neisseria gonorrhoeae* is an exclusive human pathogen that causes the sexually transmitted disease, gonorrhea. The gonococcus is highly human adapted and, thus, has developed variable mechanisms of pathogenesis that are, in part, dependent upon the site of infection (reviewed in Jerse and Rest, 1997; Dehio *et al*., 2000; Merz and So, 2000; Edwards and Apicella, 2004). In primary, human, cervical epithelial (Pex) cells, *N. gonorrhoeae* invasion results from gonococcal engagement of complement receptor type 3 (CR3) (Edwards *et. al*., 2001). In a co-operative manner, both opsonic and non-opsonic interactions occur between the gonococcus and CR3. In this regard, cervical epithelia produce a fully functional alternative pathway of complement (APC) (Edwards *et. al*., 2001), and complement (C’) is activated in response to *N. gonorrhoeae* cervical infection (Densen, 1989; Jarvis, 1994; McQuillen *et. al*., 1999; Edwards *et. al*. 2002). C’ protein C3b is deposited upon the core structure of gonococcal lipooligosaccharide (LOS) (Edwards and Apicella, 2002; Ram *et al*., 2003). *N. gonorrhoeae* surface proteins exhibit a high affinity for complement factor H (fH) binding (Ram *et al*., 1998a; Ram *et al*., 1998b; Ngampasutadol *et al*., 2008), which aids in the rapid conversion of C3b to iC3b on the gonococcal surface within minutes following gonococcal infection of Pex cells (Edwards *et al*., 2002). iC3b, bound to the gonococcus, serves as a primary ligand for CR3 adherence. However, gonococcal porin and pili also bind to the I-domain region of CR3 and are additionally required for adherence to and invasion of the host cervical epithelial cell (Edwards *et al*. 2002, Jennings *et al*., 2011).

Despite recent advances made in our understanding of gonococcal pathogenesis, factors dictating the divergent presentation of gonococcal disease observed between men and women, *in vivo*, remain unclear. That is, asymptomatic infection, generating a carrier-like state, more often occurs in *Ng*-infected women (Aledort *et al*., 2005; Densen, 1982; Densen *et al*., 1989; Bolan *et al*., 1999; Farley *et al*., 2003; Hook and Handsfield, 2008; Stupiansky *et al*., 2011; WHO, 2011). This asymptomatic state has an extreme impact on the prevalence of gonorrhea in the general population (Garnett *et al*., 1999) and in the chronic disease sequelae observed, disproportionately, among women (Bolan *et al*., 1999). Infections in women are compounded by co-infection with other transmittable organisms as well as by a number of (host) physiological factors that, cooperatively, complicate disease. For example, clinical data indicate that gonococcal pathogenesis of the female genital tract is, in part, governed by steroid hormones (SHs) (Koch, 1947; Johnson *et al*., 1969; James and Swanson, 1978; Morse and Brooks, 1985). Notwithstanding, there are limited data addressing how SHs modulate gonococcal pathogenesis. With regard to asymptomatic cervical infection, it is noteworthy that SHs exhibit anti-inflammatory as well as immunomodulatory properties (Spangler *et al*., 1969; Sonnex, 1998; Szekeres-Bartho *et al*., 2001; Watson and Gametchu, 2001; Bouman *et al*., 2005). Additionally, there are ample data to indicate that SHs modulate the activity of Akt kinase and of phospholipase D (PLD) homologs. Indeed, we show previously that under progesterone-predominant conditions reflective of the menstrual cycle luteal phase, progesterone (P4) functions in an additive manner with gonococcal PLD (NgPLD) to augment Akt kinase activity, which in turn promotes gonococcal survival within Pex cells (Edwards, 2010). Further, as CR3 and APC proteins play a critical role in gonococcal cervical disease, the ability of SHs to regulate complement production may have a profound effect on gonococcal pathogenesis in women (Sundstrom *et al*., 1989; Brown *et al*., 1990; Hasty and Lyttle, 1992; Hasty *et al*., 1993; Hasty *et al*., 1994; Norris Daju Fan *et al*., 1996; Oglesby, 1998).

Estradiol (E2) and P4 represent the major form of estrogens and progestogens, respectively, found in women. Although other estrogens and progestogens (*e. g*. estrone (E) and 17-α-hydroxyprogesterone (P2), respectively) exist *in vivo*, these hormone derivatives represent minor hormone species and, thus, are only present in minute quantities in women. *In vitro* studies examining the effect of SHs on *N. gonorrhoeae* demonstrate that some progestogens (Morse and Fitzgerald, 1974; Fitzgerald and Morse, 1976; Miller and Morse, 1977; Jerse *et al*., 2003) and estrogens (Lysko and Morse, 1980; Lysko and Morse, 1981; Salit, 1982) inhibit the growth of gonococci. However, these *in vitro* analyses were performed using estrogen and progestogen concentrations in excess of those levels reported for human blood. Hence, we set out to examine the effect(s) of physiological concentrations of E2 and P4 on *N. gonorrhoeae* as well as on complement-mediated infection of Pex cells.

## Experimental Procedures

### Cell Culture, Bacteria, and Infection Studies

Primary human cervical epithelial (*i. e*., Pex) cells were procured from surgical cervical tissue and maintained in defined keratinocyte serum-free medium (dk-SFM) (Gibco, Grand Island, NY, USA), as previously described (Edwards *et al*., 2000). Deidentified, healthy, cervical tissues were obtained from premenopausal women and were provided by the Human Tissue Resource Network/Cooperative Human Tissue Network (Columbus, OH). Pex cell monolayers were incubated in the presence of various concentrations (as noted, Fig. 1) of E2 and/or P4 for at least 24h before their use in experiments. Where applicable, hormones were maintained in the culture medium throughout the course of each assay. 17-ß-estradiol (*i. e*., E2) and progesterone (*i. e*., P4) (both from Sigma; St. Louis, MO) were selected for use in infection studies because these hormones represent the major forms of estrogens and of progestogens (respectively) found in women. To avoid potential complicating factors associated with medium containing phenol red (Moreno-Cuevas and Sirbasku, 2000), antibiotic-free medium devoid of phenol red was used in experiments designed to assess the effects of SHs on gonococcal infection. E2 and P4 were solubilized in 2-butanol (2-BtOH; Sigma) as our previous studies demonstrate that this reagent does not interfere with NgPLD activity, nor the ability of gonococci to associate with or to invade Pex cells (Edwards *et al*., 2003).

**Figure 1.**
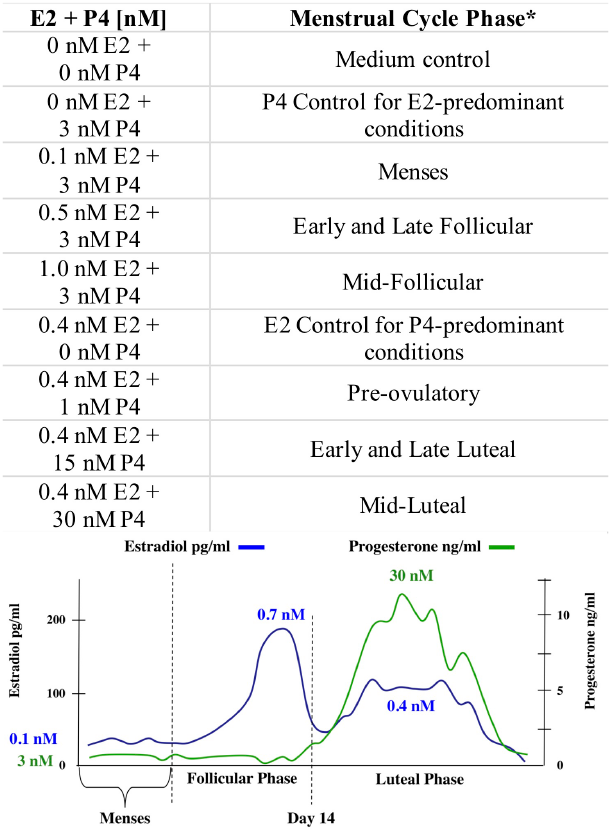
Steroid hormone concentrations used relative to the female menses cycle. During the first 14 days of the menstrual cycle (i. e. the follicular or proliferative phase), E2 levels gradually rise from approximately 0.1 nM to approximately 0.7 nM after which they again decrease around the time of ovulation. During this time, P4 levels remain relatively stable at approximately 3 nM. Within the next 14 days (the luteal or secretory phase) of the menstrual cycle, E2 levels rise slightly to approximately 0.4 nM; whereas, P4 levels rise to approximately 30 nM. The concentration of each of these hormones then decreases late in the luteal phase to approximately 0.1 nM E2 and 3 nM P4, before the onset of and during menses. * Approximate phase of the female menstrual cycle in which the combined hormone concentrations noted would be expected to occur.

*N. gonorrhoeae* strains used in the assays described herein included: 1291 (Apicella, 1974; Dudas and Apicella, 1988), 1291ΔPLD (Edwards *et al*., 2003), MS11 (Schoolnik *et al*., 1984; Segal *et al*., 1985), FA1090 (Cohen *et al*., 1994), and FA1090ΔOpa (generously provided by J. Cannon, The University of North Carolina, Chapel Hill, NC). Also examined were a panel of low-passage clinical isolates (*i. e., N. gonorrhoeae* strains IN522, SK92-679, PID-26, LT38097; all a generous gift from J. Dillard, The University of Wisconsin, Madison, WI). Strains 1291, MS11, and FA1090 were originally isolated from patients with culture-documented gonococcal disease and are commonly used to study gonococcal pathogenesis. Strains 1291 and IN522 are male urethral isolates; strain PID-26 was obtained from a patient with pelvic inflammatory disease; FA1090 and SK92-679 are genital and blood isolates, respectively, from patients with disseminated infection; and strains MS11 and LT38097 are cervical isolates obtained from women with uncomplicated gonococcal disease. *N. gonorrhoeae* strain 1291 ΔPLD contains a kanamycin-resistance cassette within the *pld* gene and, thus, does not produce NgPLD (Edwards *et al*., 2003). This kanamycin-resistance cassette lacks a promoter exhibiting functional activity in *Neisseria* and is shown to not cause polar effects in several genetic systems (Jennings et al., 1995; van der Ley et al., 1997; Power et al., 2000). FA1090ΔOpa is an unmarked deletion mutant that does not express any of the opacity associated (Opa) outer membrane proteins; these bacteria will be described elsewhere.

Infection studies were performed as previously described using a multiplicity of infection of 100 (Edwards *et al*., 2000). Pex cells were challenged with gonococci for variable lengths of time (as noted), or they were left uninfected. Uninfected Pex cell monolayers were simultaneously processed with challenged cell monolayers. Where indicated, infected and uninfected culture supernatants or Pex cell monolayers were subsequently harvested for enzyme-linked immunosorbent assay (ELISA), Western Blot, or quantitative gonococcal association and invasion analyses, as outlined below.

### Disk and Well Steroid Hormone Diffusion Assays

Estrogens (E2 and E) and progestogens (P4 and P2) were examined for their ability to inhibit *N. gonorrhoeae* growth on GC-IsoVitaleX agar plates. Estrone (*i. e*., E) and 17-α-hydroxyprogesterone (*i. e*., P2) (both from Sigma) were solubilized in 2-BtOH, as described above. An overnight agar culture of *N. gonorrhoeae* was used to inoculate Dulbecco’s modified essential medium (DMEM; Invitrogen) containing 20 mM NaHC03 (Sigma), gonococci were allowed to grow with shaking (37°C, 180 rpm), and the culture density was adjusted to 10^8^ bacterial per ml. From this liquid culture, 100 μl aliquots were removed and spread uniformly across the surface of GC-agar plates. Agar plates inoculated with gonococci were then impregnated with SHs using two methods. The first method involved the diffusion of hormones into the agar by placing vehicle control-, E-, E2-, P4-, or P2-saturated filter disks on each agar plate, *i. e*. the disk diffusion method. The second, well diffusion, method was performed essentially as described by Salit (1982). Briefly, wells were punctured within the agar surface to which hormone solutions or vehicle controls were added. Hormone solutions were allowed to diffuse into the agar before inverting the plate for overnight incubation. Concentrations of E, E2, P2, and P4 tested by either diffusion method included: 10 μg/ml (approximately 37 μM E and E2, 32 μM P4, 30 μM P2), 20 μg/ml, and 40 μg/ml; all of which are in excess of physiological concentrations; as well as 0.1 nM, 0.5 nM, or 1 nM for the estrogens or 1 nM, 15 nM, or 30 nM for the progestogens (*i. e*., physiological concentrations, Fig. 1). One percent 2-BtOH served as an assay control; however, 10% ethanol (EtOH) was also tested in that 10% EtOH was used as a solvent in similar assays described by others. Following an overnight incubation (37°C, 5% CO_2_), inhibition of *N. gonorrhoeae* growth was measured as the diameter of the area of clearing surrounding (and inclusive of) each disk or well on each agar plate, *i. e*., the zone of inhibition (ZOI). For agar plates in which a ZOI was not visible, data were recorded as the diameter of the disk (7 mm) or the well (6 mm). Disk and well diffusion assays were performed in duplicate on 4 separate occasions. Data are presented as the minimal inhibitory concentration (MIC) in which the mean ZOI exhibited a significant difference (p ≤ 0.05) in *N. gonorrhoeae* growth when compared to the 2-BtOH control. A Student’s *t*-Test (GraphPad) was used to determine the significance of data obtained.

### Far-Western Blot Analysis of the ability of steroid hormones to bind gonococcal constituents

The ability of physiological concentrations of SHs to bind gonococci was determined by Far-Western Blot analysis using methods described by Edwards *et al*. (2002). Briefly, gonococci were harvested from GC agar plates, suspended in phosphate buffered saline (PBS), and then collected by centrifugation (10,000 rpm, 5 min). The bacterial cell pellet was washed twice before resuspension in IP buffer (1X PBS, 0.5% NP-40, 0.1% SDS). 10^8^ gonococci, in sample buffer, were loaded into each well of a 4% to 12% polyacrylamide gradient gel to allow electrophoretic separation of gonococcal constituents (under denaturing conditions) and transfer to Immobilon-P membranes (Millipore Corporation, Bedford, MA). Membranes containing separated gonococcal constituents were then incubated overnight (4°C with rotation) with a median concentration (0.5 nM E2; 15 nM P2 or P4) of each hormone examined, after which the membranes were extensively rinsed. Chemiluminescent detection of hormones bound to gonococcal constituents was achieved using anti-estradiol (ESTR-1) or –progesterone (XM207) monoclonal antibodies (both from Santa Cruz Biotechnology; Santa Cruz, CA) and the appropriate peroxidase-conjugated secondary antibodies.

### Quantitative Immunoassays

SH binding to whole cell gonococci was quantitated using a modified ELISA as described by Edwards *et al*. (2002). Briefly, Pex cells were incubated with no SHs or a median concentration of E2, P2, or P4 (as noted) for 24h after which they were infected with *N. gonorrhoeae* strains 1291, MS11, or FA1090. After 30 min, the infection supernatants were harvested, and gonococci were collected by centrifugation. The bacterial cell pellet was rinsed twice with PBS, re-suspended in carbonate buffer, and 100 μl aliquots (*i. e*., 10^7^ gonococci) of each cell suspension were transferred to designated wells of a microtiter plate and allowed to dry. Gonococci-lined microtiter plates were then rinsed with PBS and blocked with ELISA blocking buffer. E2 and progestogens bound to the surface of gonococci were quantified using the anti-estradiol and –progesterone antibodies, ESTR-1 and XM207, respectively, and peroxidase-conjugated secondary antibodies. Absorbance of the o-phenylenediamine dihydrochloride peroxidase substrate was determined spectrophotometrically at 490 nm, and data were adjusted for background (no SH medium control).

The effect of SHs on the expression of Opa proteins by *N. gonorrhoeae* strains 1291, FA1090, and FA1090ΔOpa (negative control) was also determined by ELISA as described above, with slight modification. Our previous (unpublished) studies have revealed that *N. gonorrhoeae* strain FA1090 uniquely becomes cytotoxic to human, primary cervical and male urethral epithelial cells within 2 to 3 hours post-challenge. Although we have not identified the mechanism by which this strain exerts a cytotoxic effect, cell lysis does not result from a secreted FA1090 product. Therefore, to eliminate complicating factors in the data obtained as the result of the release of intracellular Pex cell constituents that could potentially modulate Opa expression (*e. g*., fatty acids, proteases), Pex cells were seeded to 24-well tissue culture dishes and allowed to grow to confluence. Pex cell monolayers were then incubated for 24h with combined variable concentrations of E2 plus P4 reflective of the female menstrual cycle (Fig. 1). A 0.2 μm pore size cell culture insert (Nalgene Nunc International, Rochester, NY) was then placed in each well to which FA1090 and FA1090ΔOpa gonococci were to be added, thereby impairing the ability of these gonococci to directly interact with the Pex cell surface while still allowing the interaction of these bacteria with secreted host products. For infections, strain 1291 was added directly to the cell monolayer, whereas the FA1090 strains were added to the cell culture insert, suspended above the cell monolayer. At 6, 24, and 48 hours post-challenge, 200 μl aliquots were removed from each well, and gonococci were collected by centrifugation. The bacterial pellet was rinsed twice with PBS, re-suspended in carbonate buffer, and culture density was adjusted to 10^8^ gonococci ml^-1^. 10^7^ gonococci, obtained from the inoculating culture as well as from infection studies, were then used to line 96-well microtiter plates, as described above. ELISAs were performed using the anti-Opa 4B12 monoclonal antibody, which recognizes a conserved domain found on all Opa proteins (Achtman *et al*., 1988).

Complement protein C3 and fH were quantified as described above for SHs and Opa expression. However, for these assays, uninfected and gonococci-infected Pex cell culture supernatants were collected and concentrated 10-fold using Centricon YM-30 (molecular weight cut off (MWCO) greater than 30 kDa) centrifugal filter units (Millipore). The filter retentates were collected and used to line 96-well microtiter plates, as outlined above. Immuno-analyses were then performed using anti-C3 (Atlantic Antibodies, Scarborough, ME; Quidel, San Diego, CA) or -fH (Quidel; Santa Cruz) antibodies.

Quantitation of the presence of CR3 on the surface of Pex cell monolayers was performed as described by Edwards *et al*. (2003). Briefly, Pex cells were passed to 96-well microtiter plates and allowed to grow to confluence. Our previous studies indicate that, during challenge of Pex cells, the *N. gonorrhoeae* mutant strain, 1291ΔPLD, is impaired in its ability to recruit CR3 to the Pex cell surface, when compared to the parental wildtype strain, 1291 (Edwards *et al*., 2003; Edwards and Apicella, 2006). Therefore, Pex cell monolayers were left unchallenged or they were challenged with wildtype 1291 or 1291ΔPLD mutant gonococci. The infection was terminated after 3h by removing the infection medium, rinsing each well thrice with PBS, and fixing the cells with 2% paraformaldehyde. Immunoassays were then performed according to standard ELISA protocols using the H5A4 anti-CD11b (the alpha subunit of CR3; The University of Iowa Hybridoma Bank, Iowa City, IA) primary antibody (Hildreth and August, 1985) and peroxidase-conjugated secondary antibodies.

All of the above assays were performed at least in triplicate and on at least three separate occasions. Negative controls for each of the above-described experiments included parallel analysis of blank wells, as well as the omission of the primary and/or the secondary antibody. Data were adjusted for background (blanks wells and/or no antibody controls), and a Student’s *t*-Test was used to determine the statistical significance of data obtained.

### Western Blot Analysis

For all experiments, polyacrylamide gel electrophoresis was performed under denaturing conditions, and western blotting was performed using methods described by Edwards *et al*. (2001). Confluent Pex cell monolayers in 35 mm tissue culture-treated dishes were incubated in various concentrations of combined E2/P4, as outlined in figure 1, 24h before infection with gonococci.

The deposition of complement proteins C3 and fH on *N. gonorrhoeae* strains 1291 and MS11 was evaluated after a 5 min infection of Pex cells. Following each Pex cell challenge, the infection supernatants were harvested, and gonococci were collected by centrifugation and rinsed twice with PBS. Total protein in bacterial cell pellets was quantified using the Coomassie Protein Assay Reagent Kit (Pierce, Rockford, IL) and adjusted to ensure equal loading. Samples were then suspended in buffer (1X PBS, 0.5% NP-40, 0.1% SDS, 40 mM iodoacetamide), pulled through a 23-gauge syringe to aid bacterial lysis, diluted 3X in protein sample buffer (final concentration: 1% SDS; 10% glycerol; 10 mM Tris-Cl, pH 7; 1 mM EDTA; 80 mM dithiothreitol; and 0.15 mg/ml bromophenol blue), boiled for 10 min, and then immediately transferred to ice. Samples were then loaded onto denaturing, 4% to 12% polyacrylamide gradient gels. Purified fH (Quidel) served as a positive control for Western Blots probed with anti-fH antibody (Quidel). As a control for C3b deposition and cleavage, gonococci were incubated (5 min and in parallel with the infections outlined above) in SH- and antibiotic-free cell culture medium spiked with normal human complement (Sigma) to a final concentration of 6%, consistent with C’ levels present at the cervical mucosa. Western Blots were probed with anti-C3 polyclonal antibody (Quidel).

### *N. gonorrhoeae* Association and Invasion Assays

Pex cell monolayers were infected with *N. gonorrhoeae* strains 1291, 1291ΔPLD, FA1090, or FA1090ΔOpa as outlined above. As indicated, Pex cells were pre-treated with E2, P4, or E2 plus P4, and hormones were maintained in (or omitted from) the culture medium over the time course assayed. Quantitative association and invasion (*i. e*., gentamicin survival) assays were then performed as we have described (Edwards *et. al*., 2000). The cytotoxicity of strain FA1090 to Pex cells prohibits their confident use in gentamicin survival assays for time periods totaling greater than 90 min. Therefore, infections were allowed to progress at 37°C and 5% CO_2_ for 1h (FA1090 strains) or 1.5h (1291 strains) without (association) or with (invasion) an additional 30 min incubation in 100 μg/ml gentamicin. The percent association with or invasion of Pex cells was determined as a function of the original inoculum and the number of colonies formed with subsequent plating of the (Pex) cell lysates. A Kruskal-Wallis analysis of variance (GraphPad) was used to determine the statistical significance of the calculated percent association or invasion for each assay. Data given are the result of 3 assays performed in triplicate.

## Results

### Physiological levels of steroid hormones are not toxic to gonococci

To begin our study, we wanted to determine the effect of physiological levels of the SHs; E2, E, P4, and P2; on the *in vitro* growth *of N. gonorrhoeae* strain 1291. SHs were tested essentially as described by Salit (1982) by both well and disk agar diffusion methods at concentrations previously demonstrated to inhibit gonococcal growth (10 μg/ ml to 40 μg/ml, the equivalent of 37 μM – 148 μM for E, 37 μM – 147 μM for E2, 30 μM – 121 μM for P2, 32 μM – 127 μM for P4) as well as physiological concentrations (0.1 nM to 1 nM for the estrogens and 1 nM to 30 nM for the progestogens; see figure 1). Physiological concentrations of SHs were not inhibitory to *N. gonorrhoeae* strain 1291; however, significant (p ≤ 0.0291) growth inhibition was observed in the presence of non-physiological levels of SHs. P4 had a greater inhibitory effect than did P2 or the estrogens, with minimal inhibitory concentrations (MICs) recorded as 40 μg/ml for E2, E, and P2 and 10 μg/ml for P4. Neither 10% EtOH or 1% 2-BtOH (controls) exerted an inhibitory effect on the *in vitro* growth of strain 1291 gonococci.

Previous data indicate that individual *N. gonorrhoeae* strains differ in their sensitivity to P4 (Morse and Fitzgerald, 1974; Lysko and Morse, 1980; Salit, 1982). Therefore, we repeated the well diffusion growth assays, described above, using a panel of *N. gonorrhoeae* strains that are commonly used to study gonococcal pathogenesis as well as several low-passage clinically isolated strains. In that transparent, Opa^-^, gonococci become increasing prevalent in women during the luteal (P4-predominant) phase of the female menstrual cycle (James and Swanson, 1978; Morse and Brooks, 1985), an Opa deletion mutant, strain FA1090ΔOpa, was also examined. Comparable to the data obtained for strain 1291, none of the gonococcal strains examined were impaired in their ability to grow on agar plates in the presence of physiological levels of E2, E, P4, or P2 (Table 1). However, the growth of individual *N. gonorrhoeae* strains did exhibit some variability in the presence of non-physiological concentrations of both estrogens and progestogens (Table 1), wherein the “clinical” isolates exhibited an increased sensitivity to E2 when compared to the laboratory strains. However, this did not appear to be associated with variable, selective, differences among strains that might be associated with the (original) anatomical site of isolation. For example, E2 exerted a greater inhibitory effect on strains IN522 and LT39087 (low-passage clinical isolates from the male urethra and the female cervix, respectively) than on strains 1291 or MS11 (laboratory strains that were originally isolated from the male urethra and the female cervix, respectively). Thus, the different inhibitory effect of non-physiological concentrations of SHs on more recent isolates, versus laboratory strains, may result from adaptation to a laboratory existence. In this regard, the Opa deletion mutant, FA1090ΔOpa, exhibited a marked increased sensitivity to all of the estrogens and progestogens tested (at non-physiological concentrations) when compared to wildtype (WT) FA1090 or the other *N. gonorrhoeae* strains examined (Table 1). Regardless, the above data suggest that the concentrations of SHs encountered by gonococci during the normal course of human infection are most likely not sufficient to inhibit gonococcal growth.

**Table 1.** Minimal inhibitory concentrations of sex hormones for N. gonorrhoeae strains.

| Minimal Inhibitory Concentration (µg/ml) |  |  |  |  |  |  |  |  |
| --- | --- | --- | --- | --- | --- | --- | --- | --- |
|  | FA1090<br>WT<sup>L</sup> | FA1090<br>ΔOpa<sup>L</sup> | 1291<sup>L</sup> | MS11<sup>L</sup> | IN522* | PID26* | LT38097* | SK92-679* |
| **E2** | 40 | 10 | 40 | 40 | 20 | 10 | 10 | 20 |
| **E** | 40 | 20 | 40 | 40 | 40 | 40 | 40 | 40 |
| **P2** | 40 | 10 | 40 | 40 | 40 | 40 | 10 | 20 |
| **P4** | 10 | 10 | 10 | 10 | 10 | 10 | 10 | 10 |
| **BtOH** | ≥ 1% | ≥ 1% | ≥ 1% | ≥ 1% | ≥ 1% | ≥ 1% | ≥ 1% | ≥ 1% |
| **EtOH** | ≥ 10% | ≥ 10% | ≥ 10% | ≥ 10% | ≥ 10% | ≥ 10% | ≥ 10% | ≥ 10% |

\*Low passage isolate; L – laboratory strain

### Steroid hormones, present at physiological levels, adhere to gonococci

Previous data indicate that the inhibitory effect of P4 on *N. gonorrhoeae* growth results from binding of this hormone to both lipid and protein constituents of gonococcal cell membranes, thereby, interfering with electron transport and, thus, respiration (Morse and Fitzgerald, 1974; Miller and Morse, 1977; Lysko and Morse, 1980; Lysko and Morse 1981). E2 is similarly shown to inhibit oxygen consumption by gonococci (Lysko and Morse, 1980; Lysko and Morse, 1981), although the adherence of E2 to gonococcal membrane constituents has not been demonstrated. P4 is a more potent inhibitor of gonococcal respiration than are E2 or P2. We did not observe an inhibitory effect by estrogens or progestogens on gonococcal growth (on agar plates) when these SHs were used at physiological concentrations. In that P4 binding to gonococci is dependent upon the concentration of available hormones (Miller and Morse, 1977), the lower concentrations of estrogens and progestogens used in our assays may not have allowed for hormone binding to the gonococcal cell membrane, which could potentially account for a lack of growth inhibition. Therefore, we wanted to determine if physiological concentrations of SHs resulted in their adherence to gonococci, as is shown previously for P4 (Morse and Fitzgerald, 1974; Miller and Morse, 1977). Far-Western Blot analysis of *N. gonorrhoeae* strains 1291, MS11, and FA1090 cell lysates using a median physiological concentration of E2 (0.5 nM), P2 (15 nM), or P4 (15 nM) revealed that, although each of these hormones bound to gonococci, P2 appeared to bind more avidly than did P4 or E2 (Fig. 2A). Only minor differences were observed among the gonococcal strains examined with regard to the individual bacterial constituents that served as targets for P2, P4, or E2 binding.

**Figure 2.**
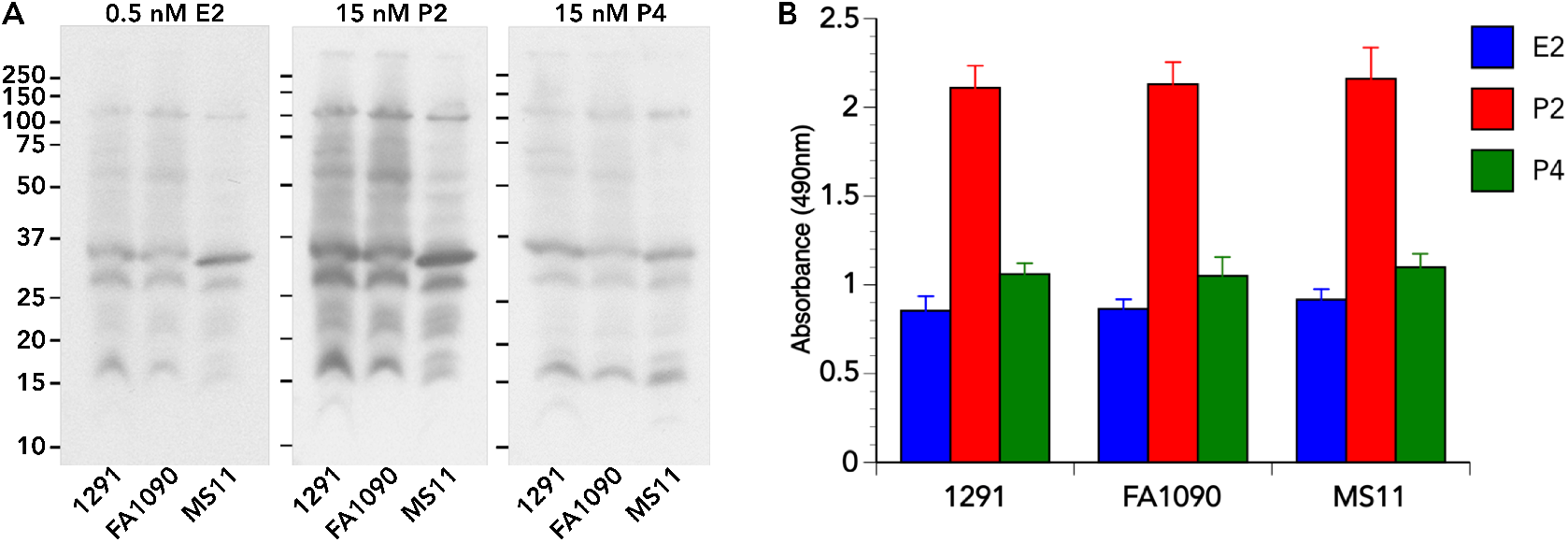
Physiological concentrations of steroid hormones bind to gonococci during Pex cell challenge. Far-Western Blot and ELISA analyses were used to examine SH binding to bacterial constituents, as described in the text. A) Gonococcal strains were loaded in triplicate onto polyacrylamide gels. Membranes were incubated with the noted concentrations of each SH and then probed with the anti-estradiol (ESTR-1) or –progesterone (XM207) antibodies, as applicable. The assay was performed on three separate occasions, yielding similar results. Representative Far-Western Blots are shown, and in which each panel was obtained from the same polyacrylamide gel/membrane. B) For ELISA analysis of SH binding to whole gonococci, *N. gonorrhoeae* were harvested from infection supernatants and used to line microtiter plates. Immuno-analysis to quantify hormone binding was performed using the anti-estradiol (ESTR-1) and –progesterone (XM207) antibodies, and data were adjusted for background (no SH medium control). There was no significant difference among the gonococcal strains examined in their ability to serve as a target for SH binding (P ≥ 0.47); however, a significant difference was observed in the ability of each SH examined to bind to gonococci (P ≤ 0.001) when compared to the no SH control. Data shown are the mean (and variance) obtained from three separate assays performed in quadruplicate.

*In vitro* data indicate that P4 binding to gonococci occurs rapidly (nearing saturation at 30 min) and in a kinetically similar manner regardless of the P4 concentration present (Miller and Morse, 1977). Therefore, to confirm our above data, as well as to determine if hormone binding to gonococci occurs under SH conditions that mimic human cervical infection, we performed an ELISA in which *N. gonorrhoeae* collected from a 30 min infection of Pex cells were used to line 96-well microtiter plates. Assays were performed using median physiological concentrations of E2, P2, or P4 (as noted above). ELISA analyses using anti-estradiol and –progesterone antibodies revealed similar data to those obtained from far-western blotting (Fig. 2B). There was no significant difference (P ≥ 0.47) among the gonococcal strains examined in their ability to serve as a target for E2, P2, or P4 binding. However, the ability of each of the individual SHs tested to differentially bind *N. gonorrhoeae* was statistically significant (P ≤ 0.001) where P2 > P4 > E2. Therefore, although P4 is a more potent inhibitor of gonococcal respiration *in vitro* than is P2 (Lysko and Morse, 1980), under the physiological SH conditions used in our assays, more P2 bound to gonococci than did P4.

### Physiological levels of steroid hormones alter Opa expression

Consistent with the increased inhibitory effect of non-physiological concentrations of SHs on transparent, Opa^-^, mutant gonococci, Salit (1982) demonstrates that gonococci allowed to grow, *in vitro*, in the presence of 1μg/ml to 15 μg/ml (or approximately 3 μM to 48 μM) P4 adapt a transparent phenotype in response to increasing P4. Although clinical data support the findings of Salit (1982), these earlier investigations relied on the use of light stereomicroscopy to evaluate Opa expression. Additionally, *in vivo*, E2 and P4 are present simultaneously in variable concentrations throughout the female menstrual cycle at ≥1000-fold lower concentrations than those used by Salit (1982). It is currently not clear if other cellular constituents alternatively, or additionally, exert selective pressure contributing to the differential expression of Opa proteins by *N. gonorrhoeae* throughout the female menstrual cycle. As the direct effect of physiological levels of SHs on Opa expression has not been examined, we quantified Opa expression by gonococci over a time course of incubation in the presence of combined variable concentrations of E2 and P4 (E2/P4) reflective of distinct points within the female menstrual cycle (see table 1).

ELISAs were performed as described in the Experimental Procedures using *N. gonorrhoeae* strains 1291, FA1090, and FA1090ΔOpa (negative control). When compared to the inocula (*i. e*., time 0), a 6 h infection of Pex cells, in the presence of E2/P4, resulted in only minor changes in Opa expression by *N. gonorrhoeae* strain 1291 (Fig. 3). Conversely, a notable increase or decrease in Opa expression occurred between 6 h and 48 h post-infection in the presence of E2/P4. A significant (P ≤ 0.001) dose-dependent increase in Opa expression by wildtype gonococci occurred by 24 h post-infection under SH conditions reflective of the follicular (E2-predominant) phase of the female menstrual cycle, which only increased modestly further in the time period from 24h to 48 h (Fig. 3A). Conversely, a dose-dependent decrease in Opa expression by the wildtype strains occurred under conditions reflective of the luteal (P4-predominant) phase of the menstrual cycle (Fig. 3B). A similar, although less pronounced, trend in Opa expression was observed when we repeated these experiments in which *N. gonorrhoeae* strains were incubated in E2/P4 in the presence or absence of Pex cells. Data corresponding to the 24h time point are shown in figure 3C and show an increase in Opa expression with increasing E2 and a decrease in Opa expression with increasing levels of P4. Only background levels of absorbance were observed in microtiter plate wells corresponding to infections performed using *N. gonorrhoeae* FA1090ΔOpa (Fig. 3C). Thus, these data are consistent with clinical data (James and Swanson, 1978; Morse and Brooks, 1985), as well as those of Salit (1982), and demonstrate that Opa expression by gonococci is responsive to SHs, which is further augmented, or inhibited, by the presence of other cellular factors produced by cervical epithelia.

**Figure 3.**
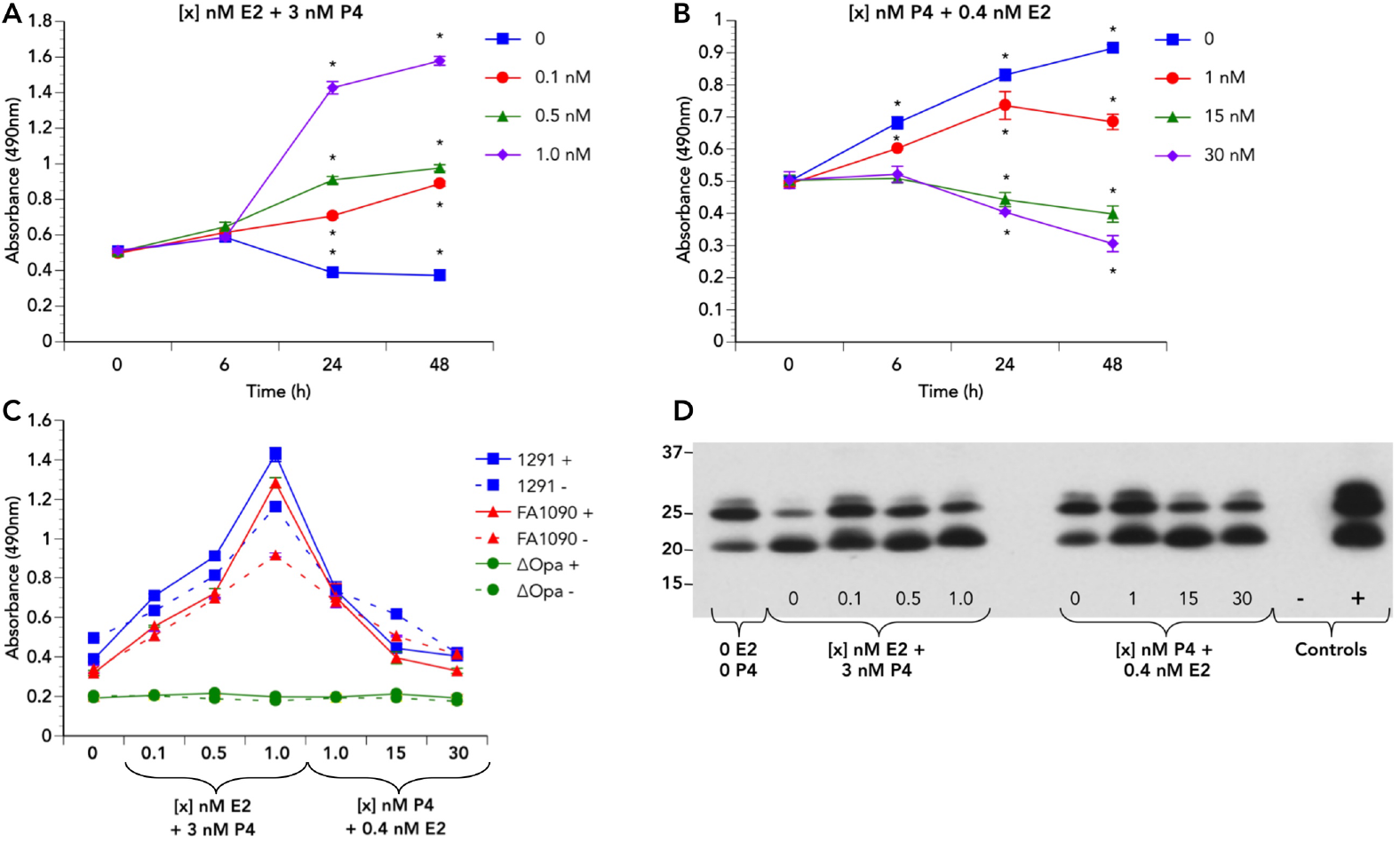
The effect of steroid hormones on Opa expression by gonococci. ELISA and Western Blot analyses were used to examine the effect of SHs on Opa expression by *N. gonorrhoeae* strains 1291, FA1090, FA1090ΔOpa, and MS11, as described in the text, and by using the anti-Opa antibody, 4B12. Data for Opa expression by *N. gonorrhoeae* strain 1291 at the noted times over a 48h time course of Pex cell challenge and under conditions reflective of the (A) follicular and (B) luteal phases of the menses cycle are shown; similar data were obtained in parallel assays performed using strain FA1090. C) Opa expression by gonococci at 24h post-incubation in E2/P4 in the presence and absence of Pex cells as it occurred for all strains examined. Data shown in panels A, B, and C are the mean (variance) obtained from a representative experiment performed in triplicate. The “0” hour time-point shown in A and B represents Opa expression measured for the inoculum used to initiate experiments. D) A representative Western Blot demonstrating the effect of E2/P4 on the Opa expression profile of *N. gonorrhoeae* MS11 is shown. Comparable data were obtained from three separate experiments. Negative control – FA1090ΔOpa cell lysate; Positive control – MS11 inoculum used to initiate experiments. * p ≤ 0.001

Under all of the E2/P4 conditions examined, *N. gonorrhoeae* strains 1291 and FA1090 expressed Opa proteins. That is, SHs did not induce a complete transparent gonococcal phenotype, as is previously suggested. Gonococci are capable of expressing zero, one, or multiple Opa isoforms, and the expression of any given Opa protein occurs independently of other Opa proteins. Thus, the expression of individual Opa proteins may be differentially influenced by the presence/absence of SHs, as is demonstrated in a mouse model of *N. gonorrhoeae* genital tract infection (Simms and Jerse, 2006). Therefore, based on the data above, we next wanted to determine the effect of variable concentrations of E2/P4 on the overall Opa expression profile of *N. gonorrhoeae* strain MS11 following a 24h challenge of Pex cells. Strain MS11 was chosen for Western Blot analysis to avoid complications associated with the use of strain FA1090 (see Experimental Procedures) and because this strain, generally, produces multiple Opa isoforms. Western blotting with the anti-Opa antibody, 4B12, revealed the expression of multiple Opa proteins by MS11 under all of the conditions assayed (Fig. 3D).

### Transparent gonococci exhibit enhanced adherence to and invasion of Pex cells in the presence of steroid hormones

Agar diffusion assays (Table 2) revealed that the FA1090ΔOpa mutant exhibited an increased sensitivity to both estrogens and progestogens at non-physiological concentrations. Although none of the bacterial strains we examined were sensitive to SHs at physiological concentrations, P4-mediated a decrease in Opa expression during Pex cell challenge, and cellular constituents produced by the mucosal epithelium could potentially alter the sensitivity of gonococci to SHs (Lysko and Morse, 1980). Therefore, to assess the effect of the P4-mediated decrease in Opa expression to cervical infection, we performed comparative *ex vivo* infection studies using *N. gonorrhoeae* strains FA1090 and FA1090ΔOpa. Pex cells were incubated with median physiological concentrations of E2/P4 (0.5 nM E2 plus 3 nM P4 or 15 nM P4 plus 0.4 nM E2) for 24 h before, as well as during, Pex cell challenge. The association of gonococci with Pex cells was significantly (p ≤ 0.0001) increased for both wildtype and mutant strains in the presence, compared to the absence, of 0.5 nM E2 (plus 3 nM P4), and this increase was significantly (p ≤ 0.0001) greater for the Opa mutant when compared to the parent, wildtype strain (Fig. 4A).

**Figure 4.**
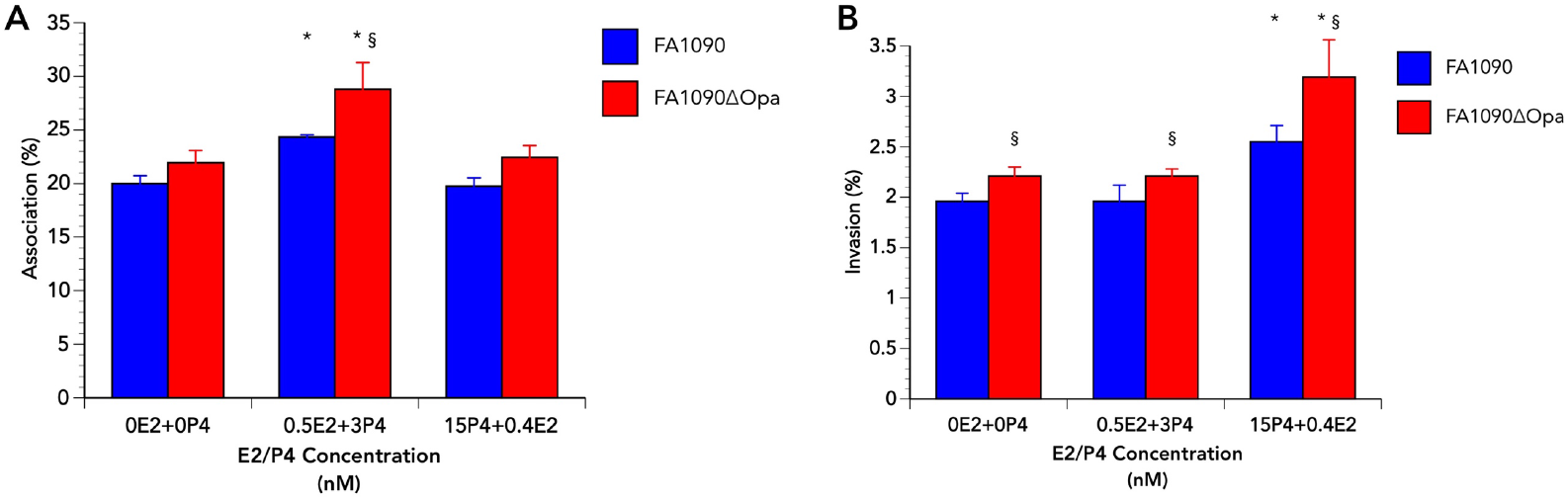
The effect of E2/P4 on the ability of *N. gonorrhoeae* strains FA1090 and FA1090ΔOpa to associate with and invade Pex cells. Association (A) and invasion (B) assays were performed as described in the text and in the presence or absence of the noted combined concentrations of E2/P4. Values given are the mean (variance) in which the percent association or invasion was determined as a function of the original inoculum and the number of colony forming units formed with subsequent plating of the cervical epithelial cell lysates. Data given were obtained from four trials performed in triplicate. *P*-values were determined using a Kruskal-Wallis k-sample analysis of variance. * p ≤ 0.0001 versus the absence of E2/P4; § p ≤ 0.0373 for comparisons between FA1090ΔOpa and wildtype FA1090.

However, 15 nM P4 (plus 0.4 nM E2) had no substantive effect on the ability of gonococci to associate with Pex cells at 1h post-infection (Fig. 4A). In the absence of either hormone, as well as in the presence of 0.5 nM E2 (plus 3 nM P4), Opa^-^ gonococci were also slightly, but significantly (P ≤ 0.0373), more invasive than wildtype bacteria (Fig. 4B). However, E2 had no significant effect on the ability of Opa^+^ (p ≥ 0.1824) *or* Opa^-^ (p ≥ 0.9999) gonococci to invade Pex cells compared to the absence of E2 (Fig. 4B). Conversely, the presence of 15 nM P4 (plus 0.4 nM E2) significantly (p ≤ 0.0001 for wildtype and mutant infections) augmented the ability of both wildtype and Opa mutant gonococci to invade Pex cells (Fig. 4B). This effect was again significantly (p ≤ 0.0001) more pronounced for the Opa mutant than it was for wildtype bacteria (Fig. 4B). These data demonstrate that gonococci of either an Opa^+^ or transparent, Opa^-^, phenotype exhibit an increased association with Pex cells in the presence of E2. Additionally, under P4-predominant conditions, the absence of Opa proteins confers a survival advantage to gonococci during Pex cell challenge, which is consistent with our previous work that indicates a beneficial effect of P4 on gonococci during Pex cell infection (Edwards, 2010).

### Estrogen augments CR3 expression by Pex cells during *N. gonorrhoeae* infection

CR3 serves as the major receptor by which gonococci adhere to and invade the cervical epithelium, *in vivo* and *ex vivo* (Edwards *et al*., 2001), by a mechanism not directly requiring Opa proteins (Edwards *et al*., 2002). Further, CR3 is recruited to the surface of Pex cells during wildtype, but not NgPLD mutant, gonococcal infection via a signaling pathway involving cervical Akt kinase (Edwards *et al*., 2003; Edwards and Apicella, 2006, Edwards, 2010). In this regard, the enhanced ability of Opa mutant gonococci to adhere to Pex cells could result from increased CR3 availability following increased surface expression by Pex cells in response to E2. If E2 and/or P4 modulate CR3 recruitment to the cervical epithelia cell surface is, currently, unknown. Therefore, we quantitated the effect of physiological concentrations of E2 and/or P4 on CR3 surface expression by wildtype- and NgPLD mutant-infected Pex cells by using a modified ELISA. These assays demonstrated that, when compared to the absence of E2, E2 significantly (p ≤ 0.021) increased CR3 surface expression on 1291 wildtype-infected cells. CR3 expression on the surface of Pex cells was highest in the presence of low (0.1 nM) E2 and (comparatively) decreased with increasing (0.5 nM and 1 nM) E2 concentrations (Fig. 5A). A similar, but less pronounced, trend in the data were obtained when uninfected and 1291ΔPLD-infected Pex cells were analyzed for CR3 surface expression (Fig. 5A). When compared to uninfected Pex cells, 1 nM P4 also induced an increase in CR3 on the Pex cell surface with wildtype infections, albeit CR3 levels did not reach those recorded for E2, and no significant (p ≥ 0.056) effect was observed when infections were performed using the NgPLD mutant (Fig. 5B). Together, these data potentially indicate that low levels of E2 trigger the recruitment of CR3 to the cervical cell surface, which is further augmented by NgPLD, when gonococci are present.

**Figure 5.**
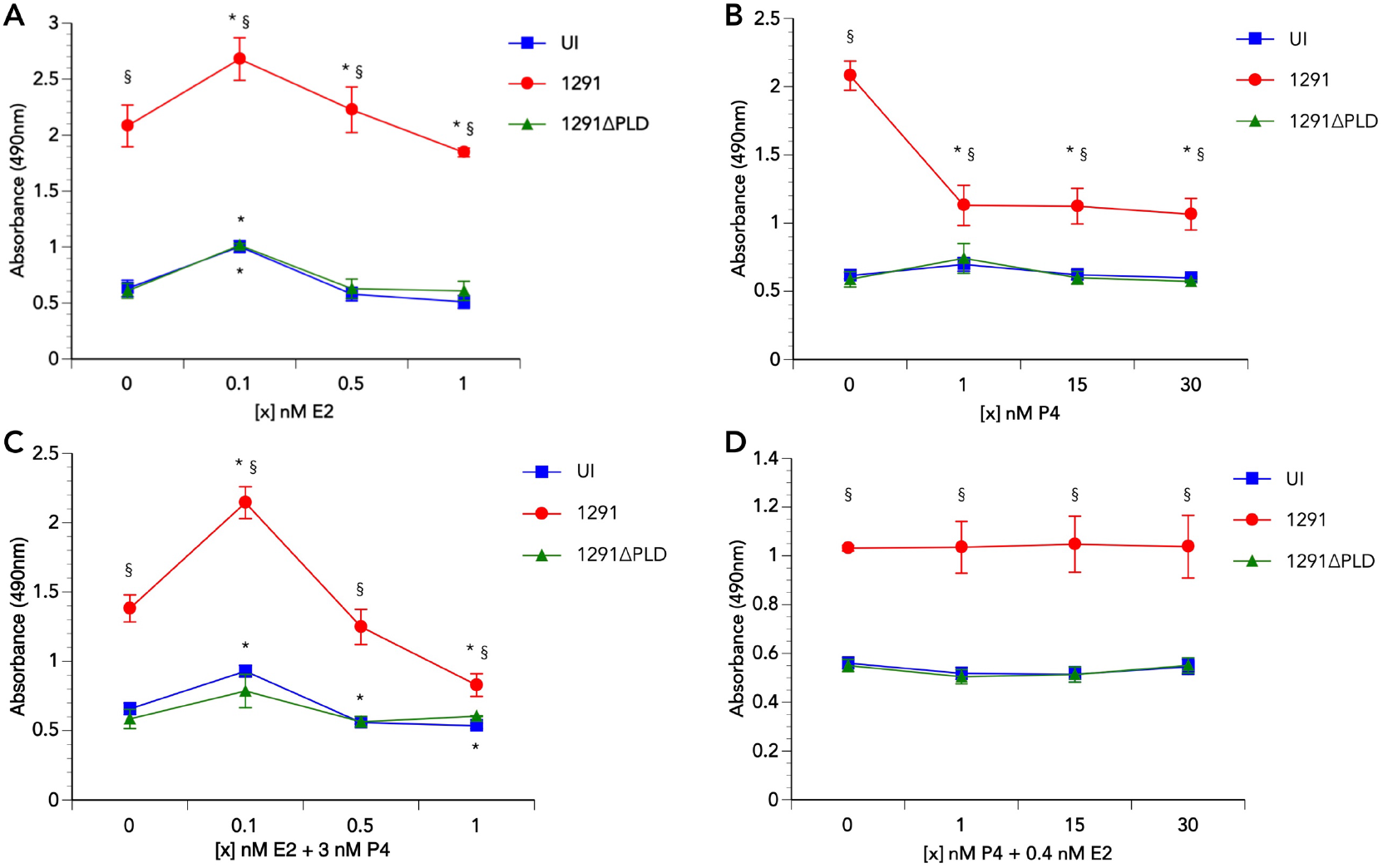
Semi-quantitative analysis of CR3 expression on the surface of primary cervical cells. A modified ELISA was used to measure CR3 on the surface of uninfected Pex cells or on cells that had been challenged for 3 h with 1291 wildtype or 1291 ΔPLD mutant gonococci, as outlined in the text and by using the monoclonal antibody, H5A4. The presence of CR3 on the cervical cell surface was measured in the presence and absence of A) E2 alone, B) P4 alone, or C and D) E2 plus P4, as noted. Values given are the mean (variance) obtained from three separate assays performed in quadruplicate. * p ≤ 0.039 for values obtained in the presence of, compared to the absence of, E2, P4, or E2/P4; § p ≤ 0.001 upon comparison to values obtained for uninfected (control) Pex cells.

We repeated our analysis under conditions more reflective of the *in vivo* environment encountered by gonococci during the follicular and luteal phases of the menstrual cycle by using combined concentrations of E2 and P4. ELISA analysis of Pex cells pre-incubated with 3 nM P4 plus various concentrations (0.1 nM, 0.5 nM, or 1 nM) of E2 (Fig. 5C), reflective of the follicular phase, yielded data similar to that observed for E2 in the absence of P4 (Fig. 5C). However, of note was that at high E2 concentrations (1 nM E2 plus 3 nM P4), the presence of CR3 on the surface of wildtype gonococci-infected cells was reduced and approached that observed for uninfected cells or Pex cells challenged with NgPLD mutant gonococci. This potentially suggests an additive effect between P4 and high levels of E2 in suppressing CR3 surface expression during *N. gonorrhoeae* infection under conditions reflective of the mid-follicular phase of the menstrual cycle (Fig. 5C, also see Fig. 1). In contrast to assays performed with P4 alone, 0.4 nM E2 plus P4 (1 nM, 15 nM, or 30 nM; reflective of the luteal phase) resulted in a modest increase in CR3 surface expression during *N. gonorrhoeae* wildtype, but not 1291 ΔPLD mutant, infection, which was generally sustained across all P4 concentrations tested (Fig. 5D). Thus, taken together, these data indicate that the effect(s) of E2, not P4, primarily augment CR3 recruitment to the Pex cell surface during *N. gonorrhoeae* infection.

To support these data, we performed quantitative association assays using Pex cells preincubated with combined SH concentrations, as noted, and that were challenged with 1291 wildtype or 1291ΔPLD mutant gonococci (Fig. 6). Under E2-predominant conditions, the association of both 1291 wildtype and 1291ΔPLD mutant (Fig. 6A) gonococci with Pex cells was increased when compared to the absence of E2, albeit a lower level of association was observed for strain 1291ΔPLD, as we have previously reported (Edwards *et al*., 2003). Gonococcal association with Pex cells was highest in the presence of 0.1 nM E2 plus 3 nM P4. Infections performed under P4-predominant conditions resulted in no significant (p ≥ 0.6682) effect on the association of wildtype *N. gonorrhoeae* with Pex cells (Fig. 6B). Although the association of 1291ΔPLD gonococci was modestly, but significantly (p ≤ 0.023), increased in the presence of 1 nM or 30 nM P4 plus 0.4 nM E2 (Fig. 6B), this increase is unlikely to be biologically relevant.

**Figure 6.**
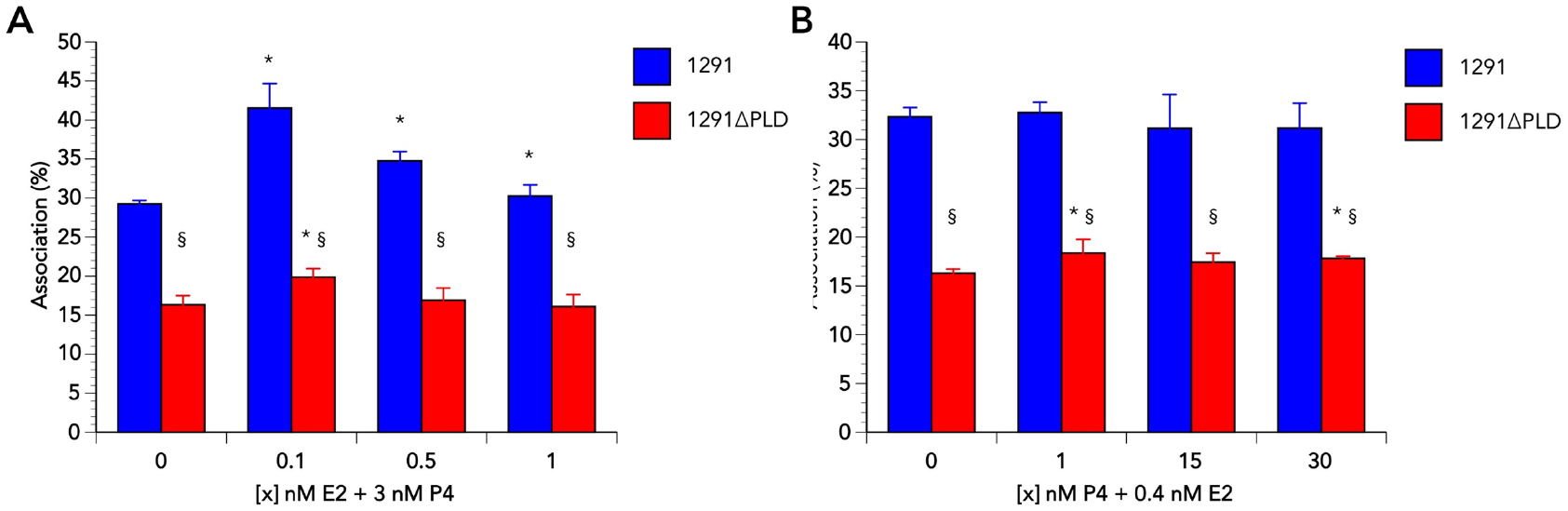
The effect of SHs on the ability *N. gonorrhoeae* 1291 and 1291ΔPLD to associate with Pex cells. The association of *N. gonorrhoeae* strains 1291 or 1291 ΔPLD with Pex cell was examined under (A) E2- or (B) P4-predominant conditions, as indicated. Values given are the mean (variance) in which the percent association was determined as a function of the original inoculum and the number of colony forming units formed with subsequent plating of the cervical epithelial cell lysates. Data were obtained from three trials performed in triplicate. * p ≤ 0.023 versus the absence of E2/P4; § p ≤ 0.0001 for comparisons between 1291ΔPLD and wildtype 1291.

These data are consistent with the ELISA data described above and support the hypothesis that under conditions in which E2 concentrations are low, *e. g*., immediately before and after, as well as during, menses, an increase in the prevalence of CR3 on the cervical cell surface results in a parallel increase in the ability of gonococci to associate with these cells.

### Complement production by Pex cells is hormonally regulated

“Inactivated” C’ protein C3, *i. e*. iC3b, is a critical opsonin required for gonococcal CR3-mediated adherence to and invasion of Pex cells. That C3 presence within the female genital tract varies in response to fluctuating hormone levels is demonstrated (Hasty, 1994; Oglesby, 1998). However, C’ production by cervical epithelia in response to SHs has not been examined, although we have shown that Pex cells produce a full alternative pathway of C’ (APC) (Edwards *et al*., 2002). In that APC proteins (*e. g*., C3, fH) play a critical role in gonococcal cervical disease (Edwards *et al*., 2002), the ability of SHs to regulate the local production of C’ could have a profound effect on gonococcal pathogenesis in women. Thereby, we examined the effect(s) of SHs on the production of the complement proteins, C3 and fH, by Pex cells. Comparative analyses of culture supernatants harvested from uninfected and *N. gonorrhoeae* strain 1291-infected Pex cells demonstrated that C’ production by cervical epithelia was responsive to E2 and P4 and, further, that infection with gonococci, not unexpectedly, augments this response (Fig. 7). Upon comparison to the no hormone control, a modest increase in C3 was observed under conditions reflective of the follicular phase of the menstrual cycle (*i. e*., 3 nM P4 plus 0.1 nM to 1 nM E2) in culture supernatants collected from both uninfected (P ≤ 0.04) and gonococci-infected (P ≤ 0.006) Pex cells (Fig. 7A). A more pronounced increase in C3 production was observed under conditions reflective of the P4-predominant, luteal phase, and a dose-dependent increase in C3 production was observed in response to increasing P4 (Fig. 7B).

**Figure 7.**
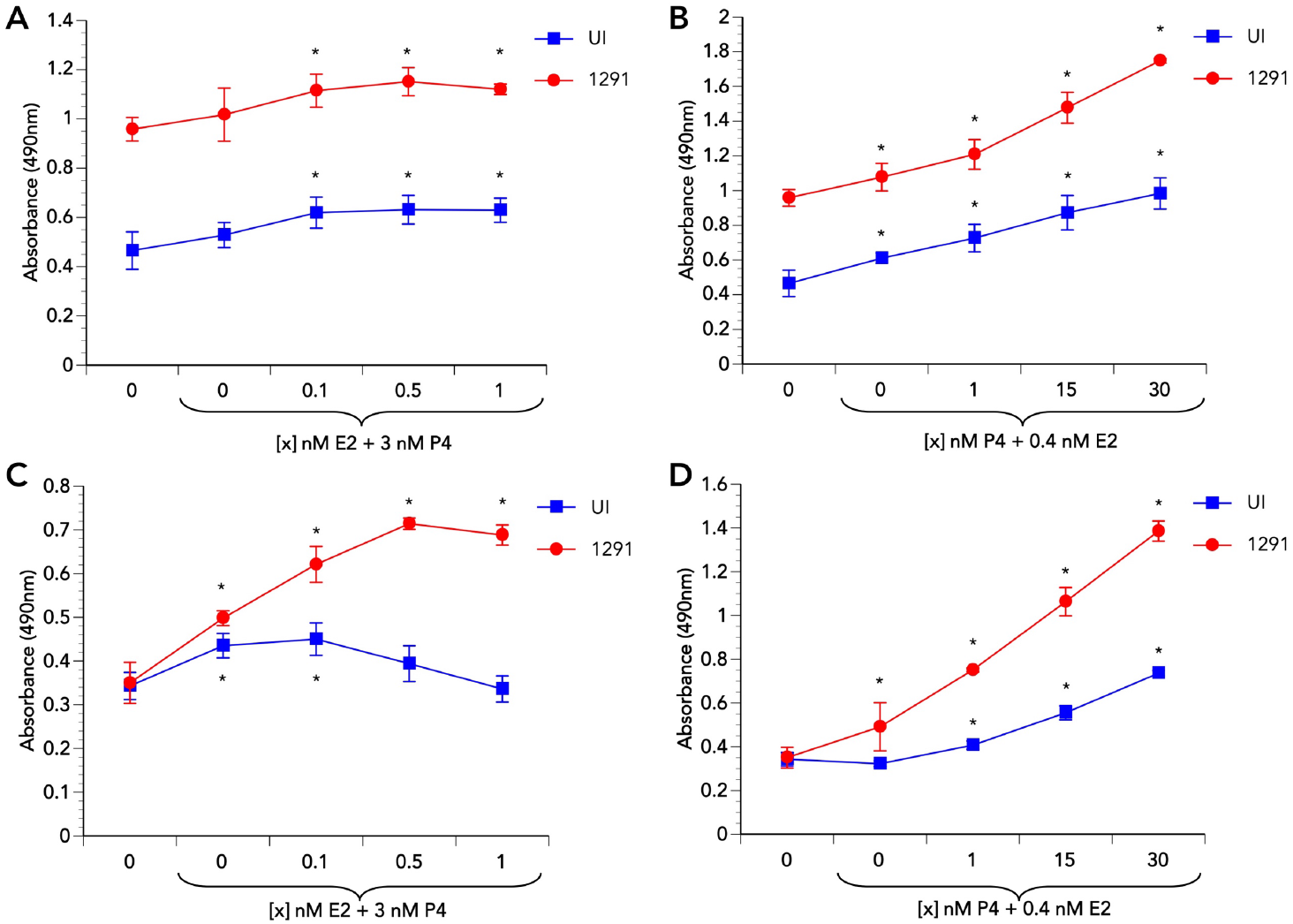
Semi-quantitative analysi s of the effect of SH s on complement, C3 and fH, production by Pex cells. Pex cells were incubated for 24h with SH concentrations reflective of the follicular (panels A and C) or luteal (panels B and D) stages of the female menstrual cycle. Pex cells were left uninfected or they were challenged for an additional 24 h with *N. gonorrhoeae* strain 1291. ELISA analysis of C3 (panels A and B) and fH (panels C and D) was performed on infected (cleared of gonococci) and uninfected culture supernatants, as described in the text, using anti-C3 or fH antibodies, respectively. Data shown are the mean obtained from three assays performed in triplicate. * p ≤ 0.04 for values obtained in the presence of, compared to the absence of, E2/P4.

Although 3 nM P4 alone or 3 nM P4 plus 0.1 nM E2 resulted in a significant (P ≤ 0.018) increase in fH in supernatants collected from uninfected Pex cells, a further increase in E2 resulted in a decrease in fH to a level that was not significantly (P ≥ 0.174) different from that of the no hormone control (Fig. 7C). However, in gonococci-infected Pex cells, a significant (P ≤ 0.012) increase in fH production occurred at all E2 (plus 3 nM P4) concentrations tested (Fig. 7C). Similar to the effect of P4-predominant conditions on C3 production, conditions reflective of the luteal phase of the menstrual cycle again resulted in a dose-dependent increase in fH production by both uninfected and gonococci-infected Pex cells, although this effect was much more pronounced in gonococci-infected cells (Fig. 7D). To our knowledge, these are the first data to examine hormonal regulation of the production of C’ directly by cells of the female genital tract mucosa. However, these data are supported by previous immunohistochemical analysis of endometrial tissue obtained from women during the luteal phase of the menstrual cycle that show a comparative increase in C3 (Hasty, 1993). Thus, the local production of C’ by the cervical epithelium is responsive to gonococcal infection as well as to SHs, in which C3 and fH biosynthesis, or their secretion. are increased during the P4-predominant luteal phase of the female menstrual cycle.

C’ component C3 exists as an (approximately) 200 kDa protein composed of an alpha and a beta chain associated by a disulfide linkage. C’ activation results in C3 cleavage to form C3a (9 kDa) and the 106 kDa (a) and 75 kDa (β) chains of C3b. Inactivation of the APC occurs during gonococcal cervical infection *in vivo* (Densen, 1989; Jarvis, 1994; McQuillen *et al*., 1999) and results from fI- and fH-mediated cleavage of the C3(b) alpha chain (106 kDa) to form iC3b [68 kDa (α_1_i or α’_1_) and 40 kDa (α_2_i or α’_2_) chains], which renders C3b incapable of fB adherence. The increased production of C3 and fH during P4-predominant environmental conditions might indicate that, during the luteal phase, more iC3b may be formed upon the gonococcal surface following C3b opsonization. To this end, we performed (denaturing) anti-C3 and –fH Western Blot analyses of gonococci collected from Pex cell infection culture supernatants to determine if the variable concentrations of SHs occurring throughout the female menstrual cycle could potentially modulate C’ deposition and its inactivation on gonococci (Fig. 8). Consistent with our previous data (Edwards *et al*., 2002), by 5 min post-infection of Pex cells, all of the C3b present on the gonococcal surface had been converted to iC3b (as indicated by the absence of a 106 kDa band and the presence of 75 kDa, 68 kDa, and 40 kDa bands on blots probed with an anti-C3 polyclonal antibody). An additional band of approximately 43-45 kDa was also visible under P4-predominate conditions, which most likely represents LOS covalently linked to C3(d). Thus, these data are consistent with our previous works and suggest the occurrence of both covalent and non-covalent interactions occurring between gonococci and the C3-α chain during Pex cell infection (Edwards and Apicella, 2002). Under conditions reflective of the follicular phase of the menstrual cycle, only minimal variability was observed in the amount of iC3b present on gonococci (Fig. 8A). Conversely, high concentrations of P4 resulted in increased iC3b present on gonococci (Fig. 8A). Similar data were obtained when parallel Western Blots were probed for the presence of fH on gonococci harvested from the same Pex cell infections used for anti-C3 Western Blots (Fig. 8B). These data are consistent with ELISA data shown in figure 7 and support the hypothesis that during the late luteal phase of the female menstrual cycle the increased production (or secretion) of C3 and fH by the cervical epithelium results in increased iC3b opsonization of gonococci.

**Figure 8.**
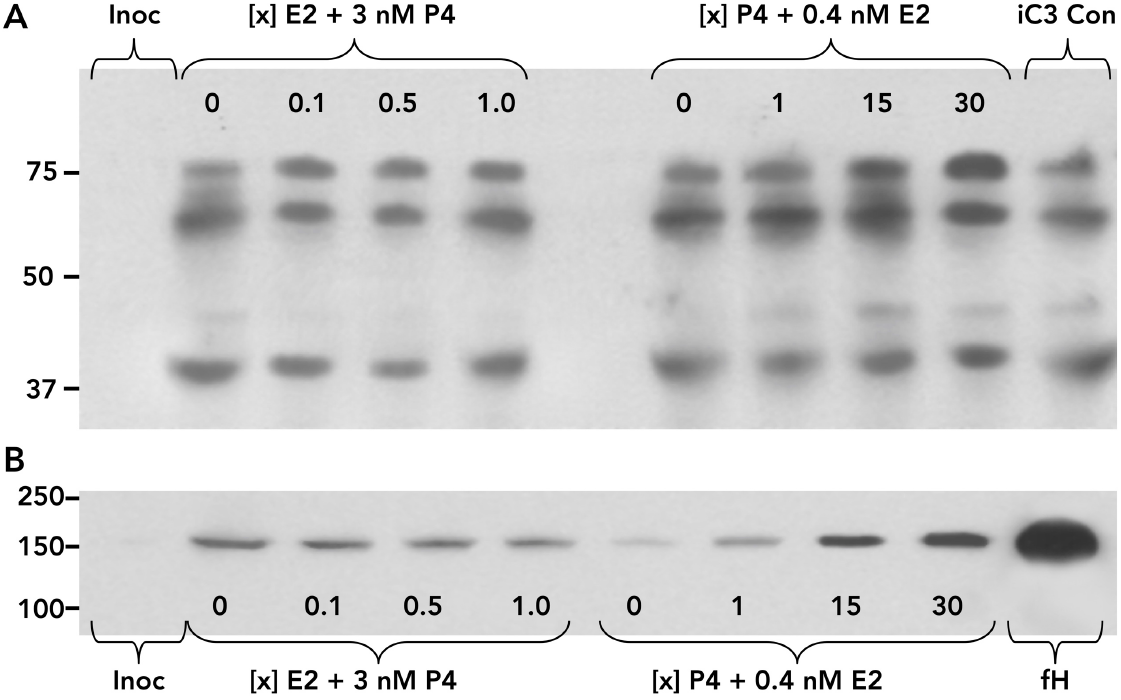
The effect of SHs on complement opsonization of gonococci during infection of primary cervical cells. Pex cells were incubated under conditions reflective the follicular ([x] nM E2 plus 3 nM P4) or luteal ([x] nM P4 plus 0.4 nM E2) phases of the female menstrual cycle at least 24 h before and during a 5 min infection with *N. gonorrhoeae* strains 1291 or MS11. Western blotting was performed as described in the text using anti-C3 (panel A) or anti-fH (panel B) antibody. A) Complete cleavage of complement protein C3 to iC3b on gonococci is demonstrated by the presence of 75 kDa (C3b-ß), 68 kDa (iC3b a_1_i), and 40 kDa (iC3b a_2_i) bands. B) The presence of fH on gonococci is indicated by the presence of a 150 kDa band. Representative Western Blots are shown and were obtained from an infection study performed using *N. gonorrhoeae* strain 1291. Negligible variability was observed in results obtained in assays performed using gonococcal strain MS 11 or from Western Blots obtained in replicate assays.

## Discussion

Although it has been realized for many years that hormone-induced, cyclic changes occurring at the mucosal epithelium of the female genital tract modulate gonococcal disease, how SHs influence the pathology of cervical gonorrhea remains poorly defined. Hence, as an extension of our previous work (Edwards, 2010), we examined the direct effect of SHs on *N. gonorrhoeae* and on complement-mediated infection of primary human cervical epithelial (*i. e*., Pex) cells.

The data presented herein were obtained using physiological levels of SHs, as they are recorded for human serum. Previous studies, performed in the absence of human cells, have indicated that estrogens and progestogens exert a bacteriostatic effect upon gonococci at concentrations (μM) exceeding physiological levels (Morse and Fitzgerald, 1974; Fitzgerald and Morse, 1976; Miller and Morse, 1977; Lysko and Morse, 1980; Lysko and Morse, 1981; Salit, 1982 Jerse *et al*., 2003). Our data support these earlier works, but also demonstrate that physiological concentrations (nM) of SHs were not inhibitory to gonococcal growth on agar plates or during Pex cell infection studies. Clinical studies demonstrate that, although gonococci are most frequently isolated from women during and around the time of menstruation, when E2 and P4 levels are low, progestin-based contraceptives increase the susceptibility of a woman to develop gonococcal disease (Fernandez *et al*., 2001; Louv *et al*., 1989; Morrison *et al*., 2004). Thus, although some disparity exists between local SH concentrations and those reported for serum, taken together, our data support the hypothesis that the actual concentrations of SHs at the level of the human cervix likely do not reach the micromolar concentrations required to inhibit gonococcal growth.

The inhibitory effect of SHs on *N. gonorrhoeae in vitro* growth results from hormone binding to the bacterial membrane, thereby, impairing oxygen uptake (Morse and Fitzgerald, 1974; Miller and Morse, 1977; Lysko and Morse,1980; Lysko and Morse 1981). However, considerable variability exists in the effect of SHs on oxygen consumption and is dependent upon environmental conditions as well as strain-specific differences observed among gonococci (Lysko and Morse, 1980). For example, acquisition of the multiple transferable resistance (Mtr) efflux system confers resistance to SH-mediated inhibition of gonococcal oxygen consumption (Lysko and Morse, 1981) and, more recently, is shown to confer a survival advantage to gonococci in a mouse model of genital tract infection (Jerse *et al*., 2003; Warner *et al*., 2007). Despite the finding that SHs inhibit *in vitro* gonococcal oxygen consumption, Fitzgerald and Morse (1976) demonstrate that P4 extends the survival of *N. gonorrhoeae* following their inoculation into chick embryos. Consistent with these data, we show that, at concentrations up to 100 nM, P4 *augments* the ability of gonococci to invade and/or survive within Pex cells, in part by augmenting host nitric oxide production (Edwards, 2010). Data presented herein also show that physiological concentrations of SHs bound to gonococcal cell lysates and to whole cell bacteria collected from Pex cell infection supernatants. One explanation for this apparent discrepancy may relate to the ability of gonococci to use nitrite and/or nitric oxide as an alternative respiratory mechanism during infection (Knapp and Clark, 1984). That is, although speculative, aerobic denitrification by gonococci might exist as an adaptation that these bacteria have evolved to promote their survival during the transition to a microaerobic/anaerobic lifestyle while in residence within the female genital tract where the binding of SHs, in particular progestogens, to gonococci would impede oxygen uptake.

Progestogen binding to the gonococcal outer membrane may play an additional role in, directly or indirectly, protecting these bacteria from proteolysis. For example, in human red blood cells, P4 binding to membrane proteins is proposed to induce a conformational change, thereby making them less susceptible to proteolytic cleavage (Ciavatti *et al*., 1974). The luteal phase of the menstrual cycle is characterized as a time of increased protein secretion by the cervical epithelium, and protease activity within the cervix increases during this period (Elstein *et al*., 1973; Profet, 1993). Several studies indicate that Opa-expressing gonococci are more susceptible to proteolysis, and clinical data indicate that transparent, Opa^-^, gonococci predominate within the female genital tract during menses and the luteal phase of the menstrual cycle (James and Swanson, 1978; Morse and Brooks, 1985). Similarly, Salit (1982) demonstrates that Opa-expressing gonococci adapt a transparent phenotype when allowed to grow, *in vitro*, in the presence of increasing concentrations of P4. Consistent with these data, we have demonstrated that E2-predominate conditions stimulate, whereas P4-predominant conditions inhibit, Opa expression by *N. gonorrhoeae* during Pex cell challenge. However, this effect was less pronounced in the absence of Pex cells, indicating that although E2/P4 directly modulate Opa expression by gonococci, other cellular factors produced by the cervical epithelium also contribute to Opa expression/suppression *in vivo*.

The *N. gonorrhoeae* proteins, porin and Opa, can serve as targets for C’ protein C3 deposition upon the gonococcal surface when they are in human serum, and C3 deposition upon these outer membrane proteins results in iC3b formation (Lewis *et al*., 2008). Whereas the gonococcus-CR3 interaction requires iC3b opsonization as well as the non-opsonic binding of pilus and porin to the I-domain region of CR3, C3 deposition occurs upon the lipid A core region of LOS, not on porin, and the direct adherence of Opa proteins to this receptor also is not required (Edwards and Apicella, 2002; Edwards *et al*., 2002). The interaction of gonococci with C’ during cervical infection likely differs from that observed in human serum in that: 1) classical complement pathway components are generally limited within the lower female genital tract [*i. e*., C4 is only detected in a small sub-population of luteal-phase cervical secretions (Vanderpuye *et al*., 1992), and we have been unable to detected C4 production by Pex cells (unpublished data)] and 2) complement is present at the mucosal epithelium at concentrations approximately 10-fold less than that observed for serum (Price and Boettcher, 1979). We have not observed a direct interaction occurring between Opa proteins and C’ protein C3 during challenge of Pex cells, which produce a fully functional APC. However, if Opa proteins do serve as a C3 target during *in vivo* mucosal infection, it then could be speculated that the decreased presence of Opa on the gonococcal surface during the menstrual cycle luteal phase, when C3 and fH production are increased, could potentially augment iC3b formation on lipid A and, thus, the gonococcus-CR3 interaction. In support of this idea, we have shown that the ability of gonococci to adhere to and invade Pex cells is enhanced when Opa proteins are not expressed. Further, we have demonstrated that during Pex cell infections performed under conditions reflective of the luteal phase of the menstrual cycle, Opa expression by gonococci is decreased, fH and C3 production is increased, and more fH as well as iC3b were observed bound to the gonococcal surface. However, a 106 kDa band, indicative of C3b, was similarly absent in anti-C3 Western Blots of gonococci isolated from Pex cell infections performed under conditions reflective of the follicular phase of the menstrual cycle (when Opa expression is not suppressed); indicating that under E2-predominant conditions all of the C3b present on gonococci also had been inactivated to form iC3b. As Opa proteins readily bind the oligosaccharide of LOS and, thereby, promote the intercellular aggregation of gonococci, an alternative explanation may be that P4-mediated suppression of Opa protein expression may render the surface of gonococci more accessible to C3 and fH, although C’ opsonization and iC3b formation would still be possible under conditions when Opa expression was not suppressed. Of note is that, although ELISA analyses demonstrated a decrease in fH production by uninfected Pex cells under conditions reflective of the late follicular phase of the menstrual cycle, fH production by gonococci-infected Pex cells increased with increasing E2 concentrations. As fH mediates C3 inactivation, these data potentially suggest that infection of the cervical epithelium by *N. gonorrhoeae* alters host signaling pathways in such a way to promote the production of fH and, thus, favor iC3b formation on the gonococcus surface where it is available to serve as a ligand for adherence to CR3. In this regard, analyses of gonococci clinically isolated from women with gonococcal cervicitis demonstrate a predominance of iC3b, vs. C3b on the surface of gonococci (Densen, 1989; Jarvis, 1994; McQuillen *et al*., 1999).

Studies performed using HeLa cervical adenocarcinoma cells demonstrate increased adhesion of several bacteria to these cells in response to E2 (Sugarman and Epps, 1982; Bose and Goswami, 1986). Although we have observed a similar phenomenon during *N. gonorrhoeae* infection of primary cervical (*i. e*., Pex) cells, HeLa cells do not express CR3 (Edwards *et al*., 2001). Further, human SH receptors (*e. g*., the estrogen and progesterone receptor), which mediate the actions of SHs, exhibit aberrant expression profiles in immortal cell lines, in that they are absent in most immortal/malignant cervical cell lines as well as in low-passage immortalized primary cell cultures (Ciocca *et al*.,1989; Nonogaki, *et al*., 1990; Macinga *et al*., 1995; Fujiwara *et al*., 1997; Kanai *et al*., 1998; Nair *et al*., 2005). There are additional data to indicate that estrogen promotes *N. gonorrhoeae* adherence to vaginal epithelial cells (Forslin and Danielsson, 1980; Forslin *et al*., 1979). The mechanism by which gonococcal adherence was enhanced was not explored; however, primary vaginal epithelium also does not express CR3 (Edwards *et al*., 2001). Additionally, during normal, *in vivo*, infection, vaginal epithelial cells are not permissive for gonococcal infection (Brooks, 1985b). We have presented data demonstrating that, with regard to *N. gonorrhoeae* cervical infection, the presence of CR3 on the Pex cell surface is responsive to E2. Our data indicate that, the density of CR3 on the Pex cell surface was greatest under conditions that mimicked those found early in the menstrual cycle follicular phase after which increasing E2 resulted in a parallel decrease in CR3 surface presence. Conversely, the presence of CR3 on Pex cells under P4-predominant conditions were similar to those observed for 0.5 nM E2 plus 3 nM P4 such as would be observed during the mid-early and mid-late stages of the follicular phase. In terms of the female menstrual cycle and *in vivo* infection these data would collectively suggest a model in which during, and immediately following, menstruation the density of CR3 on the cervical mucosa is increased and thereafter tapers off to a steady-state level during the remainder of the menstrual cycle. Menstrual blood contains C’ proteins, C’ production is increased late in the luteal phase of the menstrual cycle, and during this time, an increase in iC3b is observed on the gonococcal surface, which may poise these bacteria for increased adherence to CR3 as this receptor becomes increasingly available on the cervical epithelial surface. Consistent with this model, gonococci are most frequently isolated from infected women around the time of menses, and clinical data indicate that disseminated and ascending gonococcal infections of women occur most frequently during this same time (Brooks, 1985a; Britigan *et al*., 1985; Sweet *et al*., 1986). Similarly, women are also at an increased risk for acquiring disseminated gonococcal infection during pregnancy (Holmes *et al*., 1971; Brooks, 1985a; Hook and Handsfield, 1999), a time in which C3 levels and C’ activity are increased (Gudson, 1976).

In summary, we have provided evidence to indicate that physiological levels of SHs are unlikely to be inhibitory to *N. gonorrhoeae in vivo*. E2/P4 modulate cervical infection, in part, by regulating an increased density of available CR3 on the cervical epithelium, the increased production (or secretion) of C’ proteins by the cervical epithelium, and the decreased expression of Opa proteins by the gonococcus at select times throughout the female menstrual cycle. Taken together, our data are consistent with, and support, clinical observations pertaining to gonococcal disease in women. Investigations to further elucidate the effects of SHs, as well as E2/P4-regulated cervical constituents, on gonococcal disease in women are currently on-going in our lab.

## Acknowledgements

This work was supported by National Institutes of Health, National Institute of Allergy and Infectious Diseases grant # R01 AI076398 (J.L.E.). Cervical tissue samples were obtained from the Cooperative Human Tissue Network/Human Tissue Resource Network, funded by the National Cancer Institute. Antibody H5A4 was developed by J. T. August and J. E. K. Hildreth and obtained from the Developmental Studies Hybridoma Bank developed under the auspices of NICHD and maintained by the University of Iowa, Department of Biological Sciences, Iowa City, IA 52242. We gratefully acknowledge M. A. Apicella for his generosity in sharing bacterial strains and reagents.

